# The time course of molecular acclimation to seawater in a euryhaline fish

**DOI:** 10.1101/2021.05.17.444413

**Authors:** Lucrezia C. Bonzi, Alison A. Monroe, Robert Lehmann, Michael L. Berumen, Timothy Ravasi, Celia Schunter

## Abstract

The Arabian pupfish, *Aphanius dispar*, is a euryhaline fish inhabiting both inland nearly-freshwater desert ponds and highly saline Red Sea coastal lagoons of the Arabian Peninsula. Red Sea populations have been found to receive migrants from desert ponds that are flushed out to sea during flash floods, requiring rapid acclimation to a greater than 40 ppt change in salinity. To investigate the molecular pathways of salinity acclimation during such colonization events, a Red Sea coastal lagoon and a desert pond population were sampled, with the latter exposed to a rapid increase in water salinity. Changes in branchial gene expression were investigated via genome-wide transcriptome measurements over time from 6 hours to 21 days. The two natural populations displayed basal differences in genes related to ion transport, osmoregulation and immune system functions. These mechanisms were also differentially regulated in seawater transferred fish, revealing their crucial role in long-term adaptation. Other processes were only transiently activated shortly after the salinity exposure, including cellular stress response mechanisms, such as molecular chaperone synthesis and apoptosis. Tissue remodeling processes were also identified as transient, but took place later in the timeline, suggesting their importance to long-term acclimation as they likely equip the fish with lasting adaptations to their new environment. The alterations in branchial functional pathways displayed by Arabian pupfish in response to salinity increases are diverse. These reveal a large toolkit of molecular processes important for adaptation to hyperosmolarity that allow for successful colonization to a wide variety of different habitats.

## Introduction

Salinity is one of the main abiotic factors shaping the distribution and habitat preference of fish and other aquatic taxa. Teleosts have evolved different strategies to maintain osmotic homeostasis depending on water ion concentrations. In seawater, fish drink copiously to combat water lost via osmosis, concurrently suffering passive intake of high quantities of salts. Specialized mitochondria-rich cells of the gill epithelium, called ionocytes, are then responsible for active excess ion secretion (Edwards & Marshall, 2012; D. H. Evans, Piermarini, & Choe, 2005). In freshwater, fish face ion loss and passive osmotic water intake. To compensate for the consequent dilution of their body fluids, they actively uptake ions from the surrounding medium through specialized ionocytes (Edwards & Marshall, 2012; D. H. Evans et al., 2005). Given the profound differences in osmoregulatory mechanisms in fresh versus seawater, changes in salinity represent a significant challenge for fish, and the induced osmotic stress can lead to interferences with physiological homeostasis and impairment of biological processes (Kultz, 2015). As a consequence, most teleost fishes are restricted to limited habitat salinities (stenohaline fish). Other species, termed euryhaline fish, have evolved osmoregulatory plasticity in the organs involved in maintenance of osmotic balance (Schultz & McCormick, 2012). In particular plastic modifications to the gill epithelium, the main tissue responsible for osmoregulation in fish (D. H. Evans et al., 2005), grant the ability to live in wider salinity ranges and exploit larger habitat diversity.

Extensive work has been performed to understand the molecular underpinnings responsible for gill plasticity in euryhaline species. Early studies focused on salinity-driven expression changes of specific ion transporters and osmoregulatory genes (Deane & Woo, 2004; Scott, Claiborne, Edwards, Schulte, & Wood, 2005; Scott, Richards, Forbush, Isenring, & Schulte, 2004), expanding the existing knowledge regarding branchial ion secretion and absorption mechanisms. In particular, the role of Na^+^/K^+^-ATPase (NKA), Na^+^/K^+^/Cl^-^ transporter (NKCC1), and cystic fibrosis transmembrane conductance regulator (CFTR) in marine-type ion-extruder ionocytes was extensively investigated, and found to be highly conserved in seawater adapted teleosts (Edwards & Marshall, 2012; D. H. Evans et al., 2005). In contrast, the mechanisms for ion absorption were found to be quite diverse across different species, and several ionocytes subtypes have been described in freshwater adapted fishes (Dymowska, Hwang, & Goss, 2012; Hiroi & McCormick, 2012; Hsu, Lin, Tseng, Horng, & Hwang, 2014; Hwang & Lin, 2013). Advancements in molecular and sequencing technologies have led to the discovery of additional essential pathways for osmoregulation in euryhaline fish. Transcriptomics has provided the means to identify genes and pathways involved in osmosensing and activation of signalling cascades that initiate the osmotic stress response and the acclimation processes (T. G. Evans & Somero, 2008; Fiol & Kultz, 2007; Komoroske et al., 2016; Kultz, 2012). Profound gill remodelling, observed by microscopy and immunocytochemistry studies as alterations in ionocyte morphology and abundance (Foskett, Logsdon, Turner, Machen, & Bern, 1981; Katoh & Kaneko, 2003; Uchida, Kaneko, Miyazaki, Hasegawa, & Hirano, 2000), was linked to salinity-specific ion transporter and protein *de novo* synthesis or relocation inside the cell, activation of apoptosis pathways, and modifications of the cytoskeleton, cell-cell junctions, and the extracellular matrix (Jeffries et al., 2019; Lam et al., 2014; Mundy, Jeffries, Fangue, & Connon, 2020; Whitehead, Roach, Zhang, & Galvez, 2012). Euryhaline fish are therefore able to switch between hypo- and hyper-osmoregulation when confronted with changes in environmental salinity, although rapid increases in intra- and extra-cellular ion concentrations can negatively affect structural and functional properties of tissue macromolecules, such as proteins, lipids and DNA (Burg, Ferraris, & Dmitrieva, 2007; T. G. Evans & Kultz, 2020). Markers for an evolutionary conserved cellular stress response (CSR) triggered by macromolecular damages have been identified in gene expression and transcriptomic studies in fish exposed to changes in salinity (Brennan, Galvez, & Whitehead, 2015; Tine, Bonhomme, McKenzie, & Durand, 2010; Whitehead, Zhang, Roach, & Galvez, 2013). The CSR is comprised of defence mechanisms to protect cellular components, including the expression of molecular chaperones, and to re-establish homeostasis, by accumulation of organic compatible osmolytes and osmoregulatory mechanism switch. Moreover, the replication of damaged DNA is prevented through transcription inhibition, apoptosis and cell cycle arrest (Burg et al., 2007; T. G. Evans & Kultz, 2020; Kultz, 2005). The processes of osmotic stress response, ion homeostasis restoration and tissue remodelling are energetically demanding for euryhaline fish (T. G. Evans & Kultz, 2020; Takei & Hwang, 2016; Tseng & Hwang, 2008), and differential regulation of metabolic and mitochondrial respiration pathways during salinity challenges have been reported in several studies (Brennan et al., 2015; Chen, Lui, Ip, & Lam, 2018; T. G. Evans & Somero, 2008; Nguyen, Jung, Nguyen, Hurwood, & Mather, 2016). Given the large percentages of their energy budget consumption in response to osmotic stress, compromises in the allocation of energetic resources have been hypothesized, and consequent impairments in physiological processes such as development, growth and immune response have been reported (Breuf & Payan, 2001; Komoroske et al., 2016; Makrinos & Bowden, 2016; Morgan & Iwama, 1991). While a variety of processes involved in osmotic stress and salinity acclimation in teleosts have been defined, their ecological adaptive potential in natural colonization events is still not clear, especially the exact timing of stress, acclimation and adaptation responses and mechanisms. A number of time series salinity acclimation studies have been performed (Brennan et al., 2015; T. G. Evans & Somero, 2008; Komoroske et al., 2016; Kozak, Brennan, Berdan, Fuller, & Whitehead, 2014; Scott et al., 2004; Whitehead, Roach, Zhang, & Galvez, 2011), using designated single gene or microarray expression analyses. However, RNAseq whole-genome transcriptomics combined with an appropriate model species and collection time point selection would allow for the capture of a more complete picture of both the temporal and the mechanistic aspects of the branchial salinity response in euryhaline fish during highly saline habitat colonization.

The Arabian pupfish, *Aphanius dispar* (Rüppell, 1829), is a euryhaline species member of the Cyprinodontidae family with widespread habitat ranges (Hrbek & Meyer, 2003). Its distribution encompasses areas around the Red Sea, the Persian Gulf, the Arabian Sea, and part of the south-eastern Mediterranean basin, with populations living from inland freshwater reservoirs to coastal lagoons, and hot sulphuric springs. Along the western coast of Saudi Arabia, the Arabian pupfish is found in a large variety of environmental salinities, ranging from highly saline (41 - 44 ppt) Red Sea coastal lagoons to nearly-freshwater desert oases (0.7 - 1.5 ppt) with no permanent connection to the sea. While salinity changes have been observed to influence Arabian pupfish osmotic pressure, body ion content, and gill permeability (Lotan, 1969, 1971), no differences in performance indicators, such as resting metabolic rate, swimming speed and activity level, were caused by up to 70 ppt increases in water salinity (Plaut, 2000). In a recent genetic connectivity study supported by hydrological mapping of the region, the coastal Red Sea populations have been found to receive migrants from the inland ponds as a result of sporadic flash flood events that wash individuals out to the sea (Schunter et al., 2021). These colonization events require considerable acclimation capacities in order to survive such a rapid and drastic change in environment, especially while coping with the abrupt increase in salinity. For these reasons, the Arabian pupfish represents an excellent system to investigate the mechanisms underlying the plasticity of euryhaline fish gills during salt stress events, and the processes that allow them to acclimate long-term and colonize novel habitats. In this study, nearly-freshwater adapted Arabian pupfish from inland desert ponds were transferred to highly saline water to mimic a colonization event. The acclimation course was explored over time to disentangle the short-term processes responding to acute osmotic stress from the strategies providing the fish with longer-term adjustments required for prolonged adaptation in seawater environments.

## Materials and methods

### Experimental design

To evaluate the natural acclimation of Arabian pupfish from near-freshwater to a large increase in salinity over a short period of time, adult *Aphanius dispar* from the same genetic unit (Schunter et al., 2021) were collected using a seine net from two Saudi Arabian sites, including a near-freshwater (1.45 ppt) desert pond and a highly saline Red Sea coastal lagoon (43.49 ppm; Suppl. Table 1) between November 2015 and January 2016 (Fig. 1).

**Figure 1.**
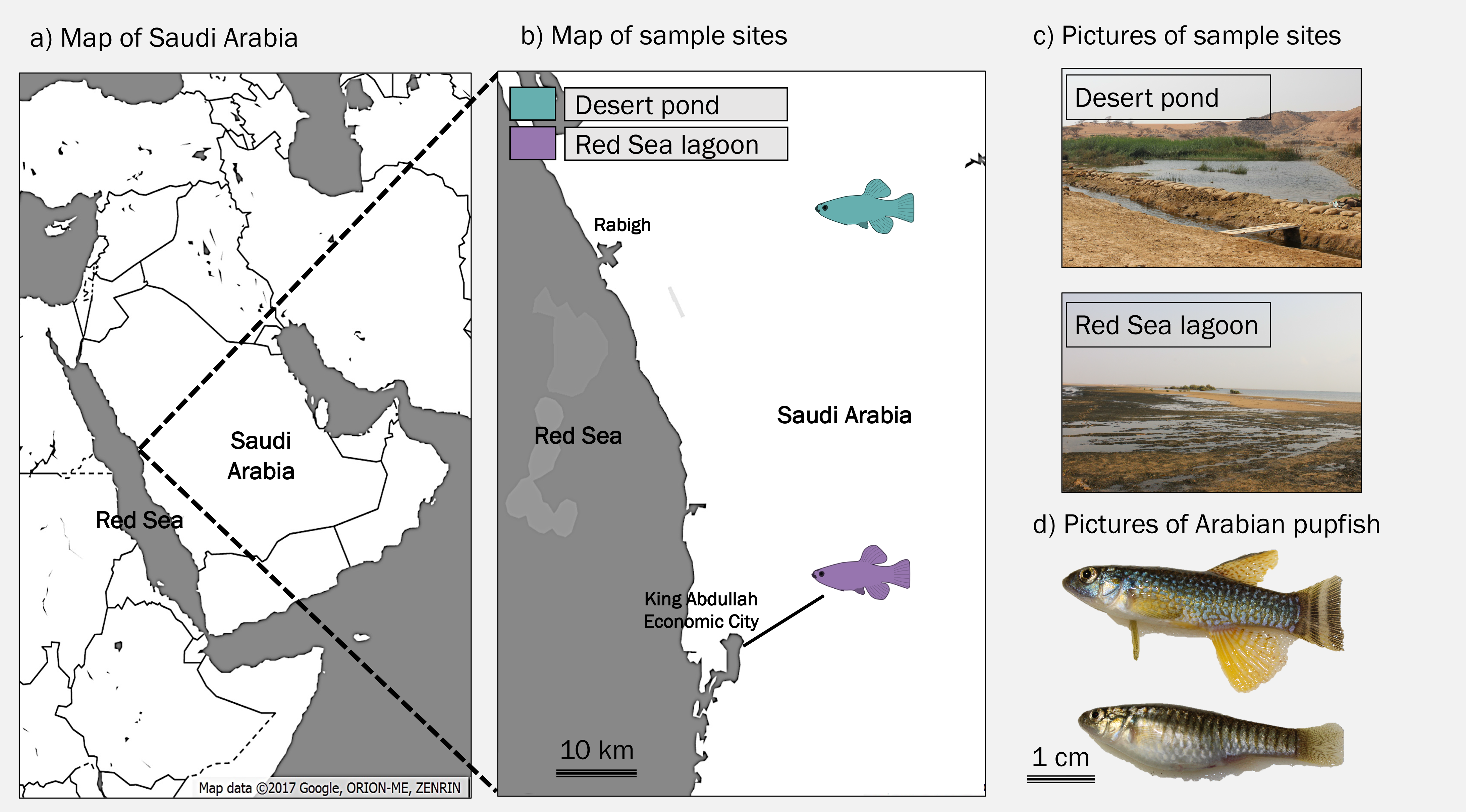
Sampling locations and target organism. (a) Map of the region with inset (b) showing the specific sampling locations in the central western region of Saudi Arabia. (c) Photos of the sampling sites. (d) Pictures of collected male (top) and female (bottom) Arabian pupfish.

The fish were transported to the King Abdullah University of Science and Technology Coastal and Marine Resources Core Lab and maintained in closed-system tanks to habituate to holding conditions for at least eight months prior to experiments. The conditions in the aquaria resembled those of the collection sites with 26°C water temperature, 8.32 ± 0.03 pH, 10L:14D photoperiod, and salinity at 1.92 ppt ± 0.01 SE for the desert pond fish and 42.68 ppt ± 0.09 SE for the coastal lagoon fish (Suppl. Table 2). Salinity was achieved using seawater, diluted for the desert pond fish with dechlorinated tap water. All fish were kept at maximum densities of seven individuals per tank, in six replicate tanks per treatment, and fed once a day *ad libitum* with commercial pelleted feed. In October 2016, the osmotic challenge experiment started: fish from the desert pond population were directly transferred to seawater, in order to mimic the change in salinity experienced during colonization of coastal lagoons of the Red Sea (Fig. 2). Temperature, photoperiod and pH were kept constant during the entire duration of the experiment in order to avoid any confounding effects. Fish were sampled pretransfer (0 h) from both populations (1.9 ppt and 43 ppt), and at five post-transfer time points (43 ppt; 6 h, 24 h, 72 h, 7 days, 21 days). At each sampling event, six individuals (one per different replicate tank) were rapidly euthanized using an overdose of tricaine methanesulphonate (MS-222, MP Biomedicals), sexed, and length was measured (Suppl. Table 3). For each individual, the right gill basket was excised and snap-frozen in liquid nitrogen and stored at -80°C for subsequent RNA extraction.

**Figure 2.**
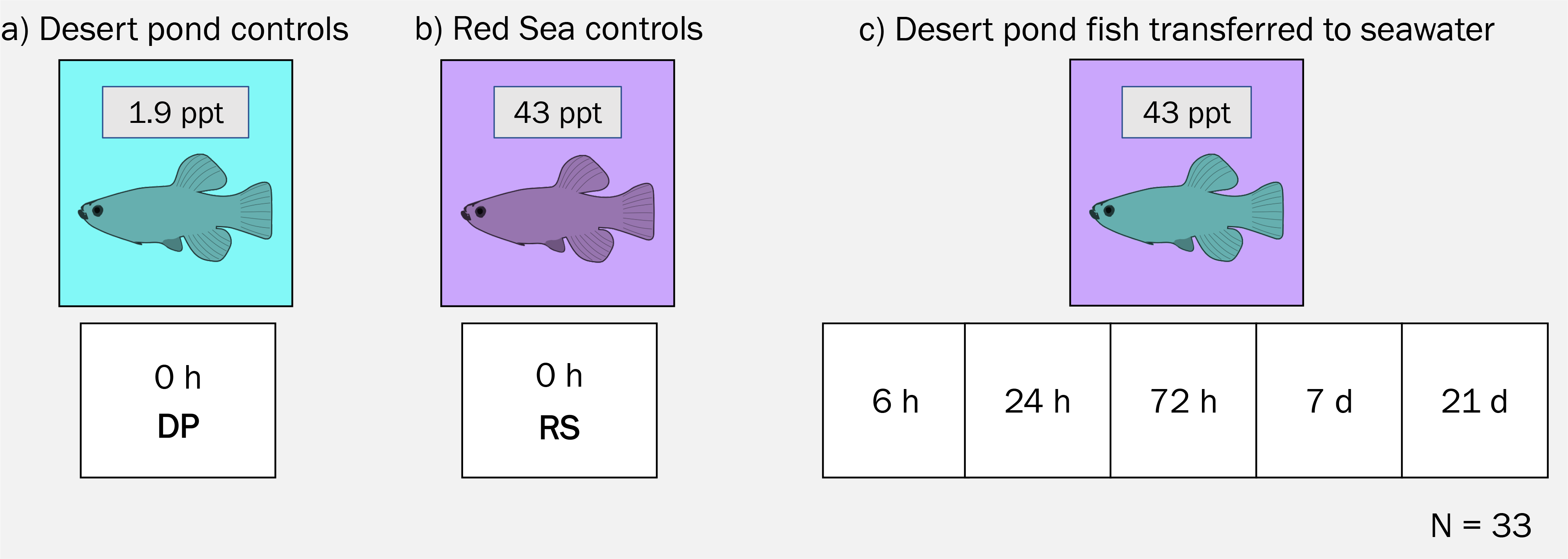
Experimental design. a) Fish from desert pond (DP) kept in native water salinity (1.9 ppt) were sampled at time 0 b) Fish from Red Sea lagoon (RS) kept in native water salinity (43 ppt) were sampled at time 0 c) Fish from desert pond transferred to seawater (43 ppt) were sampled at 5 different time points. A total of 33 samples were analyzed.

### RNA-sequencing and transcriptome assembly

RNA was isolated from gill samples using a Qiagen AllPrep DNA/RNA mini kit, following homogenization in RLT Plus buffer with MP Biomedicals FastPrep-24 homogenizer, and DNase I treated (RNase-Free DNase Set, Qiagen). Sample quality was checked using a TapeStation RNA ScreenTape assay (Agilent). Samples that did not meet RNA quality/quantity standards were excluded from the analysis. Libraries for paired-end fragments were prepared from a total of 33 samples (3-6 sample/time point) using the Illumina TruSeq stranded mRNA Library Preparation Kit, according to the manufacturer’s protocol, with each sample uniquely barcoded. Quality control check and quantification were performed with a Bioanalyzer High Sensitivity DNA assay (Agilent). Three library pools were run on an Illumina HiSeq 4000 by the King Abdullah University of Science and Technology Bioscience Core Lab.

Raw reads were processed for quality trimming and adapter removal with Trimmomatic v0.36 (Bolger, Lohse, & Usadel, 2014), using ILLUMINACLIP:2:30:10 HEADCROP:10 SLIDINGWINDOW:4:20 MINLEN:70, and inspected with FastQC (Andrews, 2010) before and after quality filtering. Trimmed reads were later corrected from random sequencing errors using a *k*-mer based method, Rcorrector (Song & Florea, 2015). Unfixable reads were discarded. Trimmed and corrected reads from all samples were used to *de novo* assemble the *A. dispar* gill transcriptome using Trinity v2.4.0 (Grabherr et al., 2011) with default settings. The assembly was then decontaminated using a hierarchical clustering algorithm called Model-based Categorical Sequence Clustering (MCSC; Lafond-Lapalme, Duceppe, Wang, Moffett, & Mimee, 2017). The MCSC pipeline was tested at five different clustering levels to determine the best one (level 3, based on the highest SS ratio), and the superclass Actinopterygii was chosen for the white list ratio calculation. Putative coding regions were predicted by TransDecoder v5.5.0 (Haas et al., 2013), and integrated with homology search against the Pfam protein domain database using HMMER v3.2.1 (Finn, Clements, & Eddy, 2011), and the UniRef90 protein database using blastp (BLAST+ v2.6.0; Camacho et al., 2009), evalue :S 1e^-5^. The decontaminated assembly was then filtered based on TransDecoder results, retaining only the single best open reading frame (ORF) per transcript. Sequence redundancy was reduced using cd-hit-est from the CD-HIT v4.8.1 package (Fu, Niu, Zhu, Wu, & Li, 2012) with 95% identity as clustering threshold. At each filtering stage, the quality of the assembly was evaluated by checking the basic alignment summary metrics, as well as quantifying the read representation by mapping the cleaned reads back to the transcriptome with Bowtie2 v2.3.4.1 (Langmead & Salzberg, 2012). Moreover, to evaluate the completeness of the assembly, and to control for potential loss of core genes during the filtering process, BUSCO v3.02 (Simao, Waterhouse, Ioannidis, Kriventseva, & Zdobnov, 2015) was run after every filtering step, using the provided Actinopterygii set of 4,584 Benchmarking Universal Single-Copy Orthologs.

### Annotation, differential gene expression and gene ontology analysis

Transcriptome annotation was performed first by BLAST searches of the TransDecoder predicted ORFs and the untranslated transcripts, using blastp and blastx algorithms respectively (evalue :S 1e^-5^), against the UniProtKb and the NCBI non-redundant (nr) databases. Where conflicts were found, the following order of priority was observed: UniProtKb/Swiss-Prot, NCBI nr, and UniProtKb/TrEMBL. BLAST results were then loaded in OmicsBox v1.4.11 (BioBam Bioinformatics, 2019), where Gene Ontology (GO), InterPro, and EggNOG (Huerta-Cepas et al., 2019) annotations were additionally performed using Blast2GO (Gotz et al., 2008).

To test for gene differential expression between the two source populations, as well as along the salinity acclimation time series, reads from each sample were first quantified using Salmon v1.1.0 (Patro, Duggal, Love, Irizarry, & Kingsford, 2017) in mapping-based mode against the *de novo* assembled transcriptome. Transcript abundance estimates were then summarized at gene level and imported in DESeq2 v1.26.0 (Love, Huber, & Anders, 2014) using the package tximport v1.14.2 (Soneson, Love, & Robinson, 2015) in R v3.6.1. Principal component analyses were run to visually check for possible overall patterns, outliers and batch effects. Three samples (DP_Rep1, T2_Rep4, T5_Rep5) were labelled as outliers and excluded from the DE analysis. Differentially expressed genes (DEGs) were identified running pairwise comparisons of control populations and post-transfer time points using the *contrast* function of DESeq2 with shrunken log2 Fold Change (log2FC) estimates by apeglm (Zhu, Ibrahim, & Love, 2019). |Log2FC| 2 0.3, False Discovery Rate (FDR) adjusted *p*-value (Benjamini & Hochberg, 1995) < 0.05 (Wald test), and a mean expression of > 10 reads (baseMean) were used as thresholds.

To identify groups of genes showing comparable trends, such as rapid or longer-term responses to the salinity challenge, DEGs revealing similar expression patterns across time points were clustered. Additionally, ImpulseDE2 v1.10.0 (Fischer, Theis, & Yosef, 2018), a Bioconductor R package specifically designed for time series data, was employed in case-only mode to discern steadily increasing or decreasing expression trajectories from transiently up- or down-regulated genes. For all clustering purposes, Red Sea population samples were treated as if they belonged to an additional time-point, the last in the acclimation timeline (long-term acclimation beyond three weeks).

Functional enrichment analyses were performed in OmicsBox v1.4.11 (BioBam Bioinformatics, 2019), with the Fisher’s Exact Test (FDR < 0.05) after removing duplicated annotations from the differentially expressed genes (DEGs) and identified gene cluster sets to find over-represented GO terms between controls and post-transfer time points.

## Results

### Transcriptome assembly and annotation

The first *de novo* transcriptome assembly for *Aphanius dispar* was created from a total of 1.38 billion raw reads with an average of 24.5 million reads per sample after trimming and error correction steps (Suppl. Table 3). These high-quality reads were used to *de novo* assemble the Arabian pupfish gill transcriptome, which resulted in 650,824 contigs with a 97.2% overall mapping rate, an N50 of 1,275 bp, and 86.2% BUSCO completeness score using the Actinopterygii database (Suppl. Table 4). A total of 2.9% contigs were filtered out in the decontamination step, and among the remaining 631,806, 141,428 contigs were predicted to contain a coding region. The final redundancy reduced transcriptome resulted in 99,167 contigs of N50 2,302 bp, E90N50 2,519 bp, with an average of 75.0% mapping rate and 86.1% of complete BUSCO genes. Tximport gene-level summarization of the final transcriptome yielded 55,451 genes. 36,863 of these genes (66.5%) were successfully annotated using SwissProt database, and another 11,078 genes had hits in the NCBI nr database. 446 more genes had positive blast hits when searched against TrEMBL, for a combined total of 48,337 annotated genes (87.2%).

### Gene expression differences in natural populations

The two sampled populations, desert pond (DP) and the highly saline Red Sea (RS) coastal lagoon, exhibited a clear branchial gene expression separation (time 0; Fig. 3), which was partly conserved throughout the salinity challenge (Suppl. Fig. 1). The pairwise comparison of fish in their original salinities resulted in 552 differentially expressed genes (DEGs; Suppl. Table 5) which is the second largest expression difference of the experiment (Fig. 4). Several functions were differentially regulated in the two populations, in particular ion transport and immune system.

**Figure 3.**
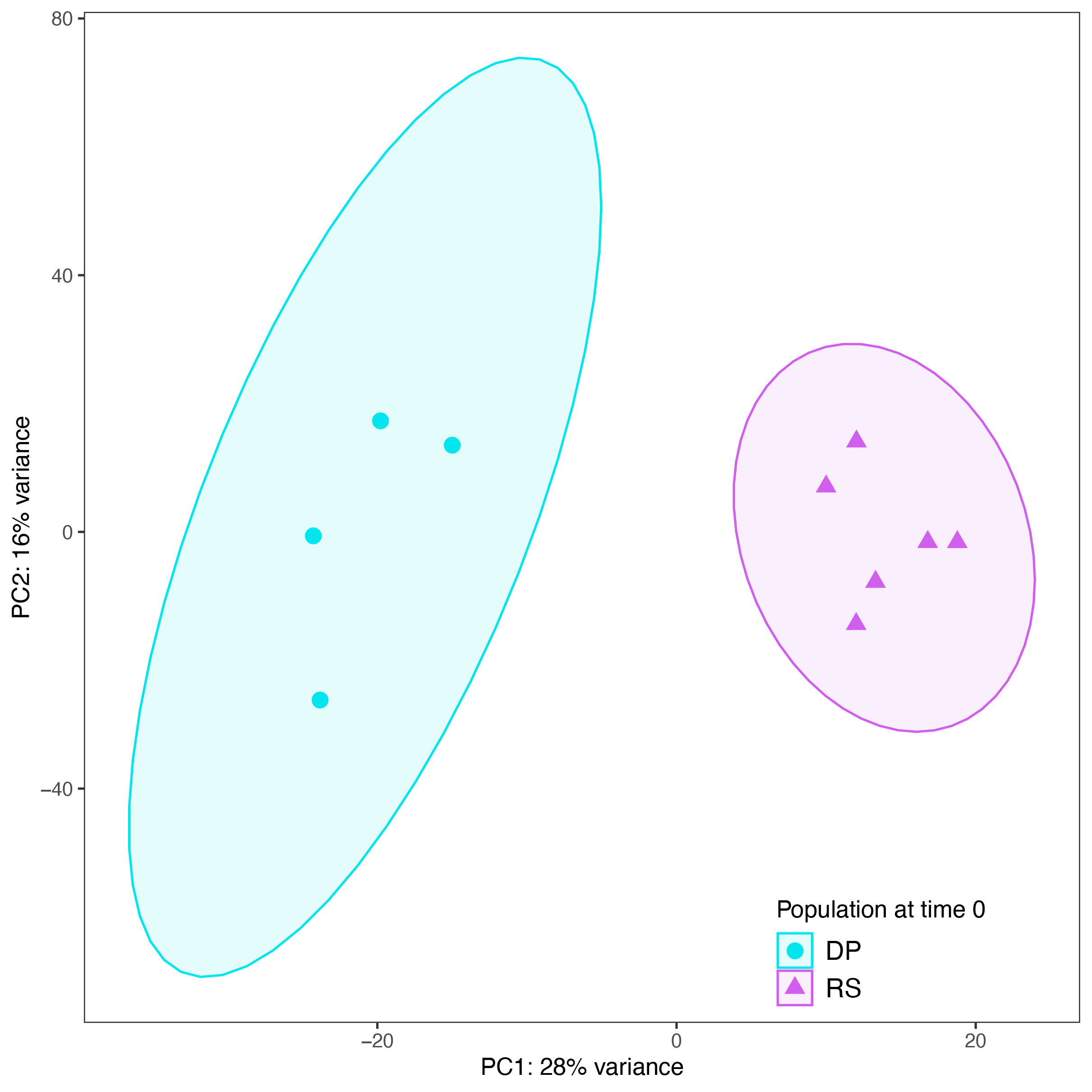
Principal component analysis (PCA) of variance stabilized expression values for the gills of desert pond (n=4) and Red Sea coastal lagoon (n=6) *Aphanius dispar* individuals at time 0. 44% of the total variation is explained by the first two components.

**Figure 4.**
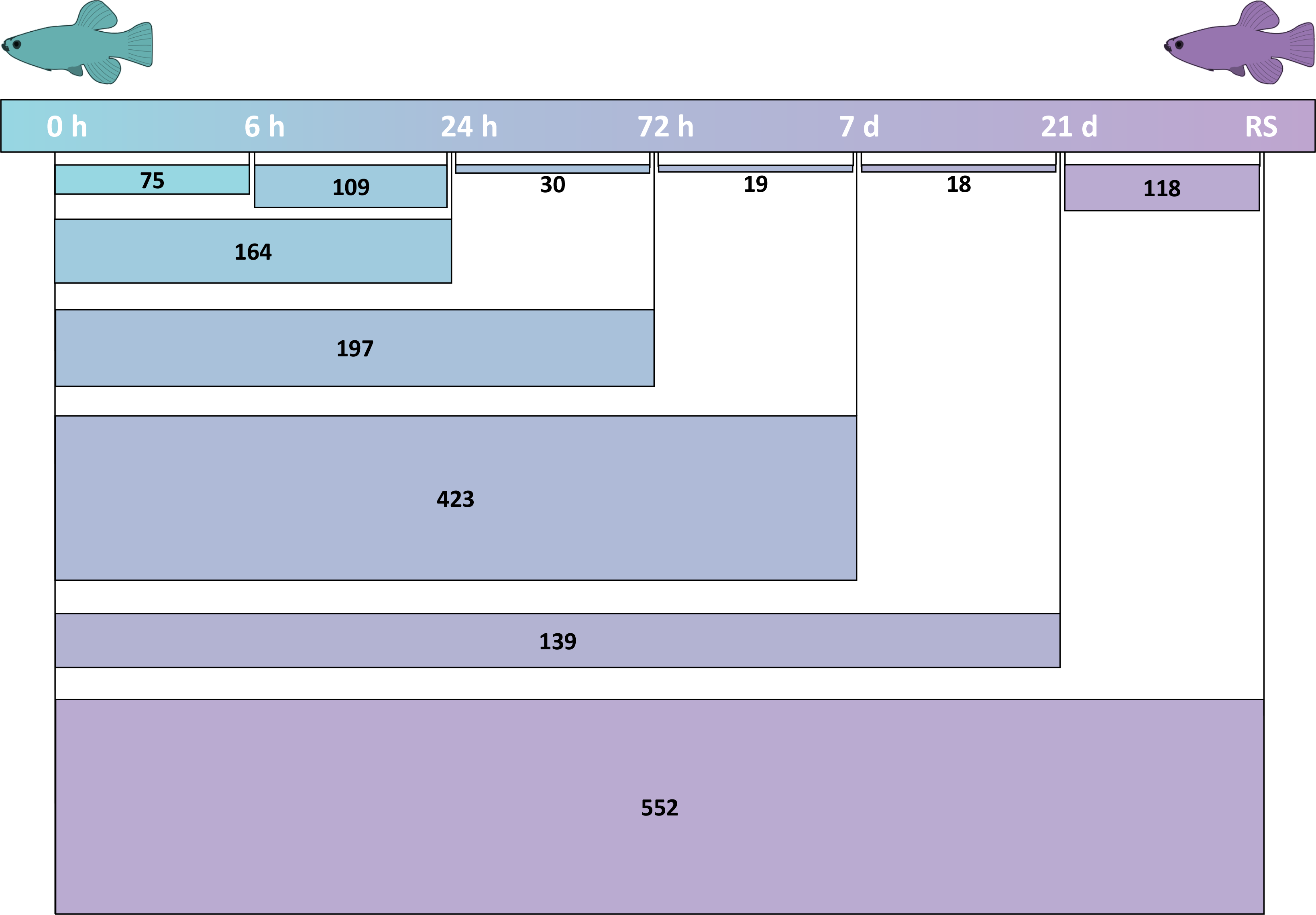
Numbers of differentially expressed genes in pairwise comparisons of desert pond fish controls (0 h) versus seawater exposed fish at different time points (6 h, 24 h, 72 h, 7 d, 21 d) and Red Sea (RS) population, and consecutive post-transfer time points.

Ion transport related terms represented 70% of total enriched GO categories between the populations at time 0 (Suppl. Table 6). The RS individuals in particular upregulated specific seawater ion transporters, such as the Na^+^/K^+^/Cl^-^ transporter (SLC12A2), the cystic fibrosis transmembrane conductance regulator (CFTR), and the transient receptor potential (TRP) cation channel subfamily V member 1 (TRPV1), together with genes involved in their regulation, such as serine/threonine kinase 39 (STK39) and the WNK lysine deficient protein kinase 2 (WNK2). Similarly upregulated in the RS population were other ion transporters and genes important in osmoregulation such as potassium channels and their modulators (KCNN3, KCNJ1, KCNJ15, ABCC8), voltage-gated calcium channel subunits (CACNA1H, CACNA1S), ammonium transporters Rh (RHAG, RHCG), sodium/hydrogen exchangers and regulators (NHEB, SLC9A2, SLC9A3R2), as well as inositol monophosphatase 1 (IMPA1) and vasoactive intestinal peptide (VIPR1). Several Na^+^/K^+^-transporting ATPase subunits (ATP1A1, ATP1A3) were also differentially expressed between the two populations, and elevated expression in the nearly-freshwater DP fish was found for a different set of ion channels, including members of the transient receptor potential (TRP) cation channel superfamily (TRPM2, TRPM5, TRPM7, TRPV4), chloride channels like the chloride channel protein 2 (CLCN2), the inward rectifier potassium channel 2 (KCNJ2), and the intracellular channel inositol 1,4,5-trisphosphate receptor type 1 (ITPR1), together with its modulator, the calcium binding protein 1 (CABP1). Aquaporin 3 (AQP3) was also upregulated in DP samples.

The two populations also showed differential regulation of the immune system. C-C motif chemokine ligands 3 and 25 (CCL3, CCL25) were upregulated in the RS population, while members of the C-X-C motif chemokine family (CXCL8, CXCL11.6, CXCL14) were upregulated in the DP fish, enriching the “chemokine activity” function. Moreover, C-C chemokine receptors (CCR4, CCR6) were upregulated in the gills of the DP population, while several genes coding for components of acquired immunity were upregulated in the seawater RS individuals, like immunoglobulins and major histocompatibility complex proteins (IGKC, IGHM, IGL1, IGKV4-1, MR1, H2-EA).

A few other functions were divergent between the populations, such as polyamine metabolism, whose related genes (ODC1, ARG1, PAOX) were upregulated in DP. Two functions were exclusive to the RS population: O-linked glycosylation, represented by five upregulated genes (B3GNT7, ST3GAL1, STR3GAL2, ST6GALNAC2, ST8SIA6), and keratinization, with the upregulation of cornifelin (CNFN), envoplakin (EVPL), keratins (XK70A, K1C1), and transglutaminase 5 (TGM5).

### Time course of the molecular responses to the salinity challenge

The selected experimental timeline and the clustering analyses of differentially expressed genes along the acclimation window (Fig. 4) allowed for the discovery of distinctively timed processes in the Arabian pupfish branchial response to the abrupt increase in salinity. While some DEGs were only transiently differentially regulated along the acclimation timeline (Fig. 5a), 231 and 269 genes steadily increased or decreased, respectively (Fig. 5b; Suppl. Table 7), and in particular the downregulated genes exhibited enrichment in cell cycle related terms (Suppl. Table 8). Through the investigation of each time point, mechanisms typical of a short-term stress response were uncovered at 6 and 24 h post-transfer, while the 72 h and 7 day time points revealed cell cycle arrest and tissue remodelling events, and the last time point (21 day post-transfer) was characterized by longer-term acclimation processes, many resembling the Red Sea population transcriptional profile (Fig. 6).

**Figure 5.**
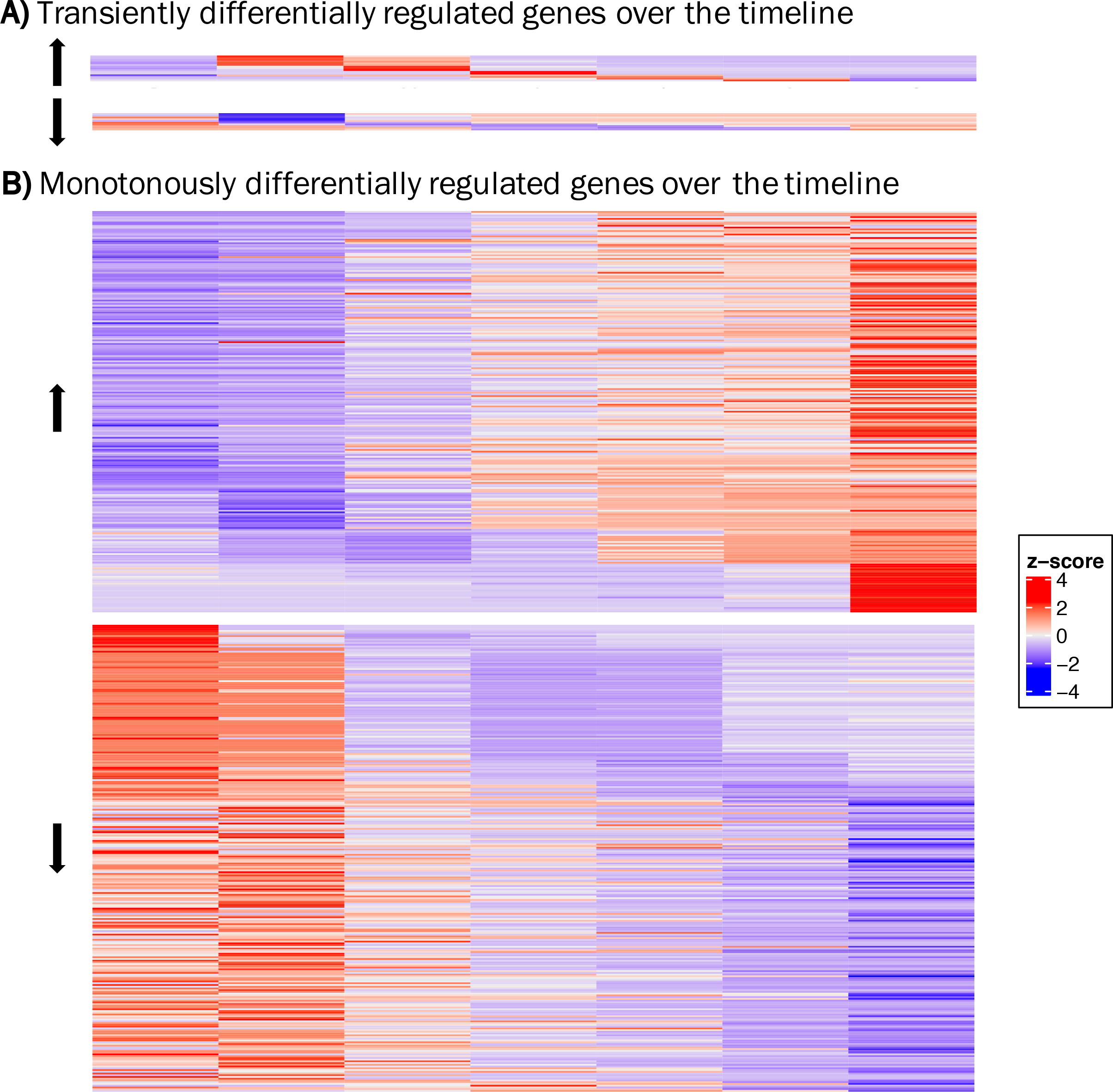
Heatmap of transiently (A) or monotonously (B) up-(i) and down- (l) regulated differentially expressed genes over the experimental timeline, as identified by ImpulseDE2 analysis. DP and RS stand for desert pond and Red Sea samples, respectively.

**Figure 6.**
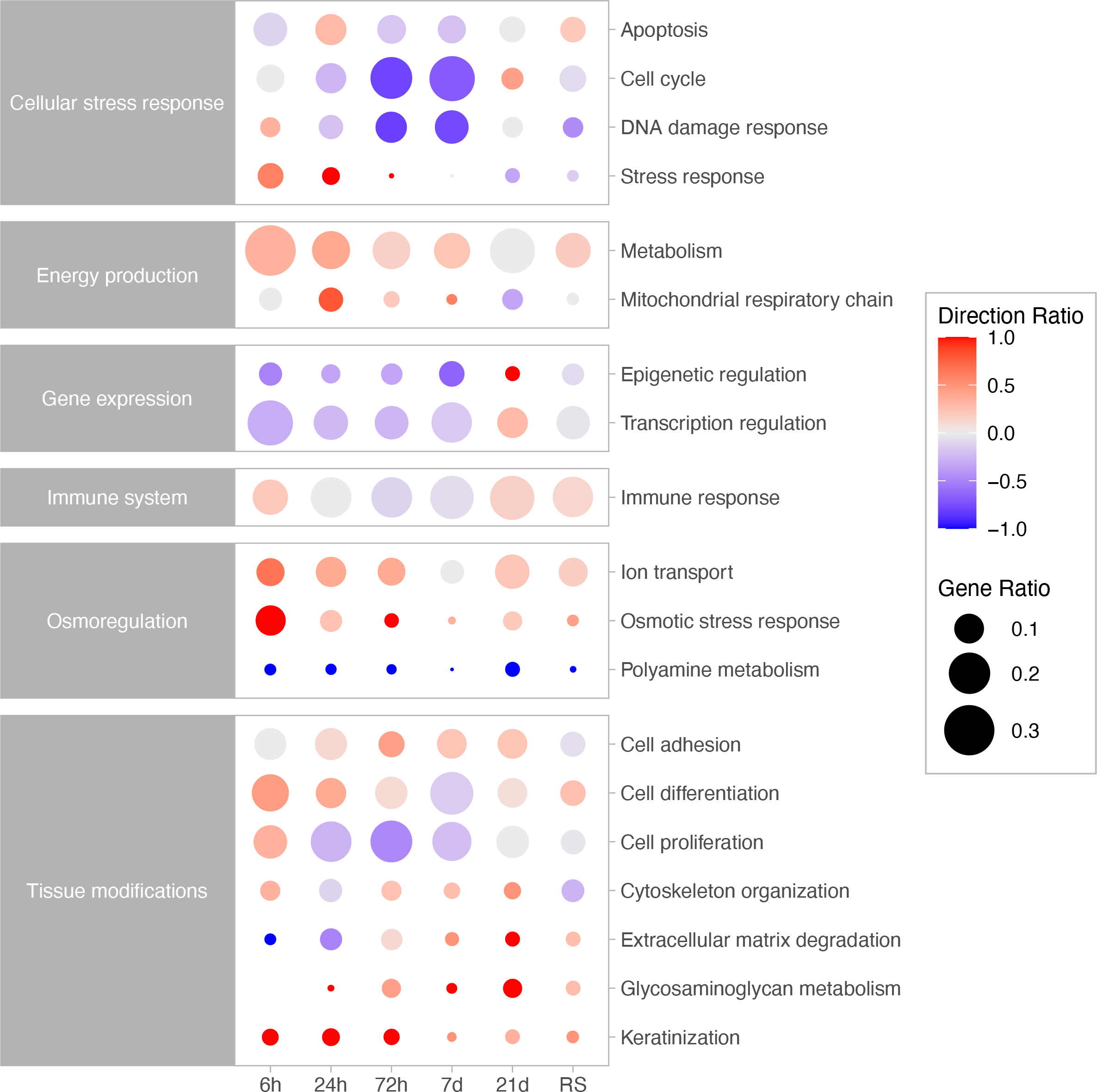
Changes in ratio and expression direction of differentially expressed genes grouped by functions along the acclimation timeline. The circle sizes are proportional to the gene number ratio for a specific function at a certain time point; the circle colours correspond to the ratio of the upregulated (red) vs downregulated (blue) genes for the function at that time point.

Just six hours after the start of the hyperosmotic challenge, 75 branchial genes showed a statistically significant change in expression compared to pre-transfer controls (Suppl. Table 9). This immediate reaction was based on a wide array of different functions (Suppl. Table 10). The acute osmotic stress caused the onset of cell signalling cascades, with the differential expression of prolactin receptor (PRLR), the stress responsive hydroxysteroid 11-beta dehydrogenase 2 (HSD11B2), which modulates intracellular cortisol levels, and serum/glucocorticoid regulated kinase 1 (SGK1), important in the cellular stress and DNA damage responses. Furthermore, the osmotic stress response was found to entail ion homeostasis pathways, through the regulation of ion transport and ion channel activity (CFTR, KCNJ2, TCAF, WNK2), and organic osmolyte synthesis and transport (GLUL, IMPA1, ISYNA1-B, SLC6A20, SLC5A7, ABCG20). In particular, ISYNA1-B was transiently upregulated between 6 and 24 hours and, in the same interval, IMPA1 showed very high levels of expression, while being upregulated across the whole timeline (Suppl. Fig. 2a). Another important function found at 6 hours post-transfer was tissue modification, by means of cellular proliferation and differentiation (GPM6B, NR4A3, NHSL1), and keratinization (CNFN, DSC1). Genes involved in the immune response were also differentially expressed at 6 hours, and class I histocompatibility antigen, F10 alpha chain (HA1F) was upregulated post-transfer. Upregulation of genes implicated in lipid (ALOXE3, ALOX15B, CPT1A) and glucose (GAD1, SLC2A8) metabolism and transport was also found. Although a few DEGs were found to be related to circadian rhythm processes (NFIL3, NR1D1, NR1D2, PER2), this result is most likely due to the sampling time occurring at a different moment of the day, rather than an effect of the salinity change.

At 24 h post-transfer, there were 164 DEGs compared to time 0 (Suppl. Table 11). “Gland development” was among the enriched GO terms (Suppl. Table 12), and included genes related to secretion, such as sodium and chloride channels (CFTR, CLCN2, SLC12A2), cell proliferation (E2F7, FGF7, FOXM1), hormonal response to stress actors (CRHR1, PRLR), cytoskeleton and extra-cellular matrix organization genes (DIAPH3, SOX9). “Lipid metabolic process” was also enriched at 24 hours, with genes related to cell membrane glycosphingolipid and glycerophospholipid biosynthesis (B4GALNT1, PLAAT4) and genes involved in lipid transport, such as carnitine palmitoyltransferase 1A (CPT1A), known to be part of the mitochondrial fatty acid oxidation pathway, and already upregulated at 6 hours. The increased energetic demand was additionally manifested by the upregulation of NADH:ubiquinone oxidoreductase complex assembly factor 4 (NDUFAF4), and solute carrier family 24 member 48 (SLC25A48), among others. Several ion transport actors and osmotic stress response regulators still played a role at 24 hours as did various transcription factors, repressors, and regulators (Suppl. Table 10). Some genes involved in DNA damage response pathway also still showed differential expression. Noteworthy, several genes involved in keratinization (CNFN, K1C1, KRT13, S100A11, TGM5) were upregulated compared to time 0, while there was a downregulation of genes involved in the extracellular matrix degradation (ADAMTS5, COL23A1, EFEMP2, PLOD2, THSD4). Tissue remodelling processes happening in transferred fish were additionally reflected by the differential expression of genes involved in the regulation of apoptosis (Suppl. Table 10), such as cytochrome C (CYC), solute carrier family 25 member 6 (SLC25A6) and voltage-dependent anion channel 2 (VDAC2), all thought to be involved in the mitochondrial apoptotic pathway.

Clustering revealed that most of the above-mentioned genes exhibited a transiently up- or down-regulated pattern in the first 24 hours, being differentially expressed compared to pre-transfer fish only in these early time points (Fig. 5a; Suppl. Fig. 2; Suppl. Table 13). The main differences between the 6 and 24 h post-transfer time points (109 DEGs; Suppl. Table 14), as portrayed by the many related enriched GO terms (Suppl. Table 15), lay in cell cycle and mitosis associated processes, with downregulation of over 30 genes with these functions at 24 hours.

Individuals from the 72 hour time point compared with the 24 h post-transfer fish resulted in only 30 DEGs (Suppl. Table 16), including the downregulation of two heat shock protein coding genes (HSPE1, DNAJC15) involved in stress response. The comparison with fish pre-transfer identified 197 DEGs (Suppl. Table 17), and the strongest signal of this contrast was the downregulation of more than 30 genes (Suppl. Table 10) involved in mitotic cell cycle and cell population proliferation (Suppl. Table 18), such for example genes fundamental for G1/S and G2/M phase transitions of the cell cycle. Concurrently, genes involved in DNA replication and repair showed downregulation compared to time 0 (Suppl. Table 10). Branchial tissue modifications were still happening at 72 hours with DEGs involved in extracellular matrix degradation including: Ca^2+^-activated cysteine protease calpains (CAPN2, CAPN5, CAPN8), and matrix metallopeptidase 8 (MMP8), known to play a role in the breakdown of the extracellular matrix during tissue remodelling, as well as several glycoproteins with cell adhesion functions, such as CEA cell adhesion molecules (CEACAM5, CEACAM6). Genes belonging to the glycosaminoglycan - or mucopolysaccharides - biosynthesis pathway were upregulated at 72 hours compared to pre-transfer conditions (Suppl. Table 10).

After 7 days from the start of the hyperosmotic challenge there were 423 DEGs compared to pre-transfer fish at time 0 (Fig. 4; Suppl. Table 19). This observed high number of differentially expressed genes could be an effect of the small sample size for this time-point (n = 3). Numerous GO terms were enriched (Suppl. Table 20), and over 80 genes, mostly downregulated compared to time 0, pertained to mitotic cell cycle as well as cell proliferation and differentiation processes (Suppl. Table 10). Seven-day post-transfer fish gills were still showing significant changes in the expression of several ion channels, as well as genes implicated in energy production and tissue modifications, such as cell adhesion, cytoskeleton organization, mucopolysaccharide metabolism, and extracellular matrix degradation (Suppl. Table 10).

The last time point at 21 days post transfer exhibited 139 DEGs (Suppl. Table 21) in comparison with pre-transfer fish. There was no functional enrichment for these genes, which were involved in a variety of processes (Suppl. Table 10). Osmoregulation was still under transition, with the upregulation of two typical seawater sodium-coupled transporters, the ATPase Na^+^/K^+^ transporting subunit alpha 1 (ATP1A1) and the solute carrier family 12 member 2 (SLC12A2). While cell cycle and energy production genes were not differentially expressed at 21 days, gene expression related genes and tissue organization were upregulated compared to time 0 (Suppl. Table 10). Moreover, two genes involved in the biosynthesis of mucopolysaccharides (GALT-1, ST3GAL1) showed upregulation, joining B3GNT7, CHST3, ST3GAL2 that were already upregulated earlier in the acclimation.

### Expression signatures leading to adaptation

Additional clustering analyses focused on the identification of genes putatively critical to long-term acclimation to elevated salinity in the Arabian pupfish. Starting from a subset of genes differentially expressed between the two populations at time 0, the focus was placed on 69 DEGs that along the experimental timeline progressively resembled the expression levels of the Red Sea population (Suppl. Table 22). Overall, some functions appeared earlier than others along the acclimation window (Fig. 6). For example, many genes implicated in osmotic stress response and osmoregulation (CA4, CFTR, CLCN2, IMPA1, KCNJ1, STK39, USP2, WNK2) were among the first to steadily change expression, between 6 and 24 hours from the start of the exposure. Cornifelin (CNFN), a component of the cornified cell envelope, and glycoprotein M6B (GPM6B), a key upstream regulator of genes involved in actin cytoskeleton organization, were upregulated at every time point post-transfer. Similarly, genes involved in the metabolism of polyamines (ODC1, PAOX), important in hyposaline acclimation in teleosts, were downregulated from the first 24 hours of exposure. As time progressed more genes changed their expression to match that of the seawater population. At 72 h post-transfer, the functions of these genes were related to changes in tissue organization and permeability, with the upregulation of calpain 2 (CAPN2) and microfibril associated protein 4 (MFAP4), both part of the extracellular matrix degradation pathway, and the downregulation of the water channel aquaporin 3 (AQP3). Moreover, some genes involved in mucopolysaccharide metabolism (B3GNT7, CHST3, GALT-1) started to be upregulated at 72 hours, while others (ST3GAL1, ST3GAL2) showed upregulation after 7 days. From 7 days of exposure to sea water, the transferred fish started to change the expression of branchial genes involved in lipid metabolism, showing upregulation of neutral cholesterol ester hydrolase 1 (NCEH1) and downregulation of low-density lipoprotein receptor (LDLR), both involved in lipoprotein and cholesterol metabolism. The transient receptor potential cation channel subfamily V member 4 (TRPV4), a non-selective cation channel involved in osmotic pressure regulation, was also downregulated from 7 day post-transfer. Finally, at the end of the experimental acclimation period, 21 days post-transfer, there was a downregulation of glycine decarboxylase (GLDC), involved in osmotic regulation, and arginase 1 (ARG1), another component of the polyamine metabolism pathway.

## Discussion

Arabian pupfish from nearly-freshwater ponds (1.5 ppt) are sporadically being flushed out to highly saline environments of the Red Sea (43 ppt) through dry riverbeds (called “wadies”) by flash floods (Schunter et al., 2021), where they are able to rapidly acclimate to the new environment and establish viable populations. Comparing the gill gene expression profiles of the native populations, as well as the changes across a salinity challenge from hours to weeks post-transfer, enabled the separation between short- and longer-term osmotic stress responses, and the investigation of the processes that allow pupfish to adapt to the high salinity typical of the Red Sea environment. The native nearly-freshwater pupfish were able to acclimate to the abrupt increase in water salinity by means of expression changes in a large number of branchial genes. A subset of genes whose expression changed with the salinity exposure to resemble those of native seawater individuals revealed the importance of related functions in long-term acclimation.

Osmoregulation was the primary function at play during pupfish acclimation to high salinity, as well as the main source of expression divergence between the two native populations. When in seawater, teleosts need to extrude passively accumulated salts through a variety of branchial ion transporters (D. H. Evans et al., 2005). Accordingly, the Arabian pupfish Red Sea population upregulated ion transporters and osmoregulation related genes typically found in marine teleosts (Hiroi & McCormick, 2012; Hwang, Lee, & Lin, 2011). The expression of many of these genes was quickly upregulated following high salinity exposure in desert pond fish and maintained along the entire acclimation timeline. For instance, genes involved in chloride ion secretion in marine-type ionocytes, such as cystic fibrosis transmembrane conductance regulator (CFTR) and Na^+^/K^+^/Cl^-^ transporter (NKCC1 or SLC12A2; Hiroi & McCormick, 2012; Marshall, 2011), were upregulated in transferred pupfish already from 6 and 24 hours, respectively. Accordingly, in another pupfish species, *Cyprinodon nevadensis amargosae*, CFTR and NKCC1 gill mRNA levels increased within a similar time frame post-transfer into seawater and remained elevated throughout the two week experiment (Lema, Carvalho, Egelston, Kelly, & McCormick, 2018), and upregulation of these two genes was also found in the gills of mummichog (*Fundulus heteroclitus*) within 24 h post-transfer from brackish to seawater (Scott et al., 2004). Involved in the activation of NKCC1 are two other genes, WNK lysine deficient protein kinase 2 (WNK2) and serine/threonine kinase 39 (STK39) (Delpire & Gagnon, 2008; Flemmer et al., 2010; Marshall, 2011), also differentially expressed across the timeline in Arabian pupfish. WNK2 has indeed been found to regulate NKCC1 activation in *Xenopus* oocytes (Rinehart et al., 2011), and its upregulation at every time point post-transfer to hyperosmotic water was also reported in the gills of the butterfish *Pampus argenteus* (Li et al., 2020). Likewise, Flemmer et al. (2010) reported the upregulation of STK39 in seawater-acclimated mummichog gills, and additionally linked it to the increased expression of NKCC1 in the same samples. The prompt upregulation of marine-type ion transporters and osmoregulatory genes in desert pond pupfish exposed to seawater to quickly resemble the Red Sea population levels confirms the importance of these genes in osmoregulation to highly saline conditions and reveals the high degree of conservation in osmoregulatory mechanisms among different fish species, Arabian pupfish included.

Another fundamental component of teleost osmoregulation in salt water is the *myo*-inositol biosynthesis (MIB) pathway, through which the compatible organic osmolyte *myo*-inositol is synthetized and accumulated inside cells during osmotic stress for protection from salinity-induced damages (Yancey, Clark, Hand, Bowlus, & Somero, 1982). The two enzymes constituting the MIB pathway, inositol monophosphatase 1 (IMPA1) and *myo*-inositol phosphate synthase (MIPS or ISYNA1), were both differentially expressed in high-salinity exposed pupfish. Accordingly, both genes were also upregulated following seawater transfer in other euryhaline fishes, such as eels (*Anguilla anguilla*; Kalujnaia, McVee, Kasciukovic, Stewart, & Cramb, 2010) and turbots (*Scophthalmus maximus*; Cui et al., 2020), where the MIB pathway knockdown was directly implicated in causing weakened gill osmoregulation and reduced survival (Ma et al., 2020). The MIB pathway is therefore an important osmoregulation mechanism in seawater across several different species. However, the transient upregulation of MIPS in the first 24 hours only highlights the importance of this pathway in the short-term osmotic stress response of the Arabian Pupfish.

In hyposaline waters fish are susceptible to passive ion loss and need to compensate by active uptake of osmolytes from the surrounding water (Edwards & Marshall, 2012). In Arabian pupfish, chloride uptake is likely accomplished by chloride ion channel protein 2 (CLCN2), typically found in freshwater ionocytes (Leguen, Le Cam, Montfort, Peron, & Fautrel, 2015; Wang, Yan, Tseng, Chen, & Hwang, 2015), which was highly expressed in desert pond individuals pre-transfer, and decreased from 24 hours of high salinity exposure onward. Rainbow trout (*Oncorhynchus mykiss)* ionocytes (Leguen et al., 2015) and a Sacramento splittail population exposed to seawater *(Pogonichthys macrolepidotus*; Mundy et al., 2020) also show this pattern, indicating the importance of this chloride channel in a variety of fish species inhabiting freshwater environments, where chloride uptake from the surrounding water is a key aspect of osmoregulation. Conversely, desert pond Arabian pupfish lack most of the previously described mechanisms for the uptake of sodium (Dymowska et al., 2012; Hsu et al., 2014), and might possibly exclusively rely on specialized isoforms of Na^+^/K^+^-ATPase (NKA) subunits for its import. Although the NKA gene is usually identified in marine acclimated teleosts, where it is part of the ion secretion machinery, in Arabian pupfish different transcripts annotated to the a-1 subunit gene were upregulated in desert pond individuals, and only one transcript was upregulated in Red Sea pupfish. While this could be an indication of population-specific NKA subunit a-1 isoforms (Mundy et al., 2020), it is likely that salinity-dependent isoforms with opposite functions of ion uptake in hyposaline water and salt secretion in seawater exist in this species, as previously described for several euryhaline teleosts (Bystriansky, Richards, Schulte, & Ballantyne, 2006; McCormick, Regish, & Christensen, 2009; Richards, Semple, Bystriansky, & Schulte, 2003; Tipsmark et al., 2011; Urbina, Schulte, Bystriansky, & Glover, 2013; Velotta et al., 2017). In the climbing perch (*Anabas testudines*) and in salmonids, NKA a-1 isoform a expression levels are highest in freshwater and decreases post-transfer to seawater, while other isoform mRNA expressions increase following exposure (Bystriansky et al., 2006; Ip et al., 2012). Accordingly, some of the NKA a-1 subunit coding transcripts showed a steady decreasing pattern along the exposure timeline in transferred pupfish, with one transcript resulting in a 4.5-fold downregulation at the end of the experimental timeline, which could be an indication of acclimation to the seawater habitat, where sodium uptake is not needed anymore. Revealed Arabian pupfish hyperosmoregulatory mechanisms could represent a new model for fish branchial ion absorption at low salinities and confirm the wide diversity of evolutionary distinct branchial adaptations to hyposaline environments in teleosts. Overall, the main mechanisms differentiating Arabian pupfish natural populations pertain to osmoregulation, and rapid acclimation from near-freshwater to highly saline waters involves adjusting the gill gene expression to resemble the osmoregulatory processes typical to the long-term adapted seawater population.

The second major difference between the two Arabian pupfish populations concerned the immune system, with several genes involved in the inflammatory and immune responses also differentially regulated post-transfer. In teleosts, salinity has been known to have intricate impacts on the immune system. While osmotic stress has been found to increase the nonspecific immune response, a depression of the acquired immune response has also been reported owing to trade-offs in resource allocation (Makrinos & Bowden, 2016). Red Sea population and translocated desert pond fish showed however overexpression of acquired immune response components, such as immunoglobulins (Ig) and major histocompatibility complex class I-related (MR1). The immune response capacities are therefore likely not impacted during acclimation to seawater in Arabian pupfish. Other fishes have been shown to not suffer from immune depression in seawater, such as Nile tilapia *(Oreochromis niloticus*; Dominguez, Takemura, Tsuchiya, & Nakamura, 2004), as well as *Acanthopagrus latus* and *Lates calcarifer*, where plasma Ig levels increase with water salinity (Mozanzadeh et al., 2021). Likewise, hyperosmotic immersion of *Paralichthys olivaceus* boosts branchial major histocompatibility complex expression and the overall mucosal immune response 24 to 48 h post-exposure (Gao, Tang, Sheng, Xing, & Zhan, 2016). Indeed, a crucial role in immunity and defence in teleosts is exerted by mucosal surfaces (Salinas, 2015). Fish gills, skin and gut are coated with a thin mucus layer which acts as a barrier from the surrounding environment and is characterized by physical and antimicrobial defensive functions (Koppang, Kvellestad, & Fischer, 2015; Reverter, Tapissier-Bontemps, Lecchini, Banaigs, & Sasal, 2018), but a role in osmoregulation has also been suggested (Shephard, 1994; Wong et al., 2017). Two mucin-like transcripts were upregulated in Red Sea Arabian pupfish, as well as at 24 h and 7 days post-transfer in desert pond fish. Moreover, several genes related to O-linked glycosylation of mucins, glycosaminoglycan metabolism and mucus production (Malachowicz, Wenne, & Burzynski, 2017) were upregulated both in Red Sea individuals and in seawater-exposed desert pond fish. Accordingly, in *Anguilla japonica* mucosal tissues, seawater elicits an increase in mucus cell numbers and secretion, possibly to trap sodium ions (Wong et al., 2017). Similar findings were reported for *Salmo salar*, in addition to salinity-driven modifications of mucin biochemistry (Roberts & Powell, 2003), that were also later described in other euryhaline fishes (Mylonas et al., 2009; Roberts & Powell, 2005). As the molecular results suggest, increased gill mucus production might be at play in seawater in Arabian pupfish and could be another aspect of their acclimation strategy to hyperosmotic environments.

While osmoregulation and immune response were revealed to be important in long-term adaptation to seawater, a series of short-term and transient response mechanisms were also elicited during the acclimation timeline. Abrupt increases in ion concentration can lead to macromolecular damages in exposed fish epithelia. An increase in sodium, for example, has been linked to cell membrane damages (T. G. Evans & Kultz, 2020) through lipid peroxidation, catalysed by lipoxygenases in response to stress (Kultz, 2005). Two lipoxygenases (ALOXE3, ALOXE15B) were indeed upregulated in Arabian pupfish gills 6 h post-transfer, and were also identified in similar seawater transfer experiments on eels (Kalujnaia et al., 2007). Such membrane disruption can lead to the activation of the so-called cellular stress response (CSR), which encompasses defence mechanisms to protect and repair damaged cellular components and restore homeostasis (T. G. Evans & Kultz, 2020; Kultz, 2005). The CSR machinery responds to sodium-destabilized proteins by overexpressing molecular chaperones, such as heat shock proteins (T. G. Evans & Kultz, 2020), two of which were temporarily upregulated in Arabian pupfish at 24 hours, as well as in a hyperosmotic challenged Sacramento splittail population (Mundy et al., 2020). Rising intracellular sodium levels can also cause nucleic acid structural disruptions (T. G. Evans & Kultz, 2020), and consequently, increased expressions of DNA damage related genes are often reported as part of the CSR following hyperosmotic stress in teleosts (Brennan et al., 2015; Su, Ma, Zhu, Liu, & Gao, 2020; Whitehead et al., 2013). In Arabian pupfish several DNA damage related genes were transiently upregulated especially in the first 24 hours of exposure, like serum/glucocorticoid regulated kinase 1 (SGK1), and similarly increased in other fish following acute seawater challenges (T. G. Evans & Somero, 2008; Shaw et al., 2008). Another aspect of hyperosmotic-induced CSR, at least in cultured human cells, is the inhibition of transcriptional and translational activities (Burg et al., 2007). Equivalent to other species, such as the climbing perch (Chen et al., 2018), Arabian pupfish exhibited a downregulation of transcription related genes following seawater exposure, which might be a mechanism to prevent the replication of high salinity-damaged DNA. An inhibition in transcription and translation might also explain the onset of cell cycle arrest that was identified in both Arabian pupfish and climbing perch (Chen et al., 2018) via a downregulation of large sets of cell cycle and mitosis involved genes during the acclimation to seawater. In support of these findings, an immunocytochemistry study in tilapia (*Oreochromis mossambicus*) observed a G2 phase arrest in the mitotic cycle of gill cells over a period of 16-72 h post-seawater exposure (Kammerer, Sardella, & Kultz, 2009). Hence, in the first hours of high salinity challenge, gill acclimation in Arabian pupfish is dominated by the onset of macromolecular damages followed by the cellular stress response machinery initiating the repair of the compromised processes to restore homeostasis. Days to weeks after the start of the hyperosmotic challenge, an inhibition of transcriptional activity and simultaneous cell cycle arrest might represent a strategy for the fish to prevent the replication of damaged DNA, preserve energy and buy time to respond to macromolecular damages caused by the increase in ion concentration.

For longer-term acclimation to seawater, euryhaline fish gills must undergo profound remodelling events in order to switch from an ion absorbing epithelium to an ion secreting one. Arabian pupfish started displaying processes involved in tissue remodelling from the first hours of salinity exposure. Likely to allow a rapid reorganization of the gill epithelium to reverse the ion transport direction, a transient increase in cell proliferation and differentiation related genes was uncovered, as seen in the gills of euryhaline tilapia in the first eight hours of seawater exposure (Kammerer et al., 2009). At the same time, genes involved in keratinization, a process by which keratin accumulate inside epithelial tissue cells to provide barrier-like functions, started to be upregulated. Keratinization gene expression has been found to be salinity dependent in tilapia (Ronkin, Seroussi, Nitzan, Doron-Faigenboim, & Cnaani, 2015) and to be upregulated following air exposure in the amphibious mangrove rivulus skin (*Kryptolebias marmoratus*; Dong et al., 2021). Keratinization may represent a strategy to reduce the amount of water loss during dehydration, possibly also following increased environmental salinity, as seen in Arabian pupfish. Programmed cell death, or apoptosis, of high salinity-damaged cells and freshwater-type ionocytes represents another of the first steps in gill epithelium remodelling, essential for full acclimation to seawater (T. G. Evans & Kultz, 2020). A transient upregulation of cytochrome c (CYC) and other genes involved in the mitochondrial apoptotic pathway was seen in Arabian pupfish at 24 hours. This has been previously recorded in mummichog in response to osmotic stresses (Whitehead et al., 2012), and is supported by a microscopy study in Mozambique tilapia revealing increased branchial apoptotic freshwater ionocytes one day after transfer to seawater (Inokuchi & Kaneko, 2012). At 24 h post-transfer, genes related to cell adhesion began to be upregulated, and cytoskeleton and extracellular matrix organization functions showed increased expression from 72 hours of exposure. Cell adhesion and extracellular matrix pathway upregulation was similarly reported in Sacramento splittails between one and seven days into the acclimation to elevated salinity (Jeffries et al., 2019; Mundy et al., 2020), and branchial cell cytoskeleton reorganization is largely recognized as a fundamental aspect of salinity acclimation in teleosts (T. G. Evans & Somero, 2008; Fiol & Kultz, 2007; Nguyen et al., 2016). As a consequence of tissue remodelling events, an upregulation of mitochondrial respiratory chain and metabolism related genes is also expected to support the increased energy demand, as previously found in similar experiments of euryhaline fish translocation (Chen et al., 2018; Lam et al., 2014; Whitehead et al., 2012). Analogously, in Arabian pupfish there was an overall upregulation of metabolism related genes up to 7 days post-transfer, while genes involved in mitochondrial respiration were overexpressed especially between days one and seven, which is consistent with the time frame for major gill remodelling processes in other species (Foskett et al., 1981; Katoh & Kaneko, 2003; Mundy et al., 2020). In a similar fashion to other euryhaline teleosts, seawater exposed pupfish are therefore affected by transient and longer-lasting gill tissue modifications occurring from the first hours to several days after the beginning of the exposure, and potentially resulting in perdurable modifications which allow longer-term acclimation to the highly saline environment.

Arabian pupfish inhabit profoundly divergent environments of the Arabian Peninsula, ranging from nearly-freshwater ponds found in desert areas to highly saline Red Sea coastal lagoons. The plasticity of these fish under steep increases in water salinity plays a major role in the colonization potential of this species. Remarkably, Arabian pupfish are able to survive flash flood events which likely displace them from desert oases and wash them to the sea, where they eventually establish viable populations (Schunter et al., 2021). By simulating and analysing the exposure to high salinity from near-freshwater over time, not only key processes for a successful acclimation were identified, but also the importance of their timing was uncovered. Arabian pupfish branchial salinity-elicited pathways revealed osmoregulation, immune system and mucus production to be rapid but also long-term acclimation mechanisms to the new environment. In the short-term, cellular stress response processes were triggered, which prevented the fish from suffering permanent damages following acute hyperosmotic exposure. Later in the acclimation, pathways involved in gill epithelium modification and remodelling equipped the organism with lasting adaptations to the increased salinity. While some of the processes occurring during the acclimation timeline resembled mechanisms of seawater exposure previously reported in other euryhaline fish species, others, such as increased mucus production and keratinization, represent less common strategies for high salinity acclimation in teleosts. Overall, the branchial processes revealed in this nearly-freshwater Arabian pupfish population during high salinity acclimation sheds light into this non-model euryhaline species colonization potential of seawater habitats. A large set of differentially timed molecular mechanisms plays a role in the plastic reorganization of the gills in hyperosmotic environments that allows for the expansion of euryhaline teleosts into a wide variety of different habitats.

## Acknowledgements

This study was supported by the King Abdullah University of Science and Technology (KAUST). The project was completed under KAUST ethics permit 15IBEC35_Ravasi. We thank KAUST Coastal and Marine Resources Core Lab and Jessica L. Norstog for assistance with animal collection and maintenance. We also thank KAUST Bioscience Core Lab for assistance with Illumina library preparation and sequencing. The authors declare there are no conflicts of interest.

## Author contributions

L.C.B., A.A.M., C.S. conceived, designed and performed the experiment, with input from T.R. L.C.B. and C.S. performed the sample collection. L.C.B. and A.A.M. prepared the samples for RNA sequencing. L.C.B. analyzed the data, with input from C.S. and R.L. L.C.B. interpreted the data and wrote the manuscript, with input from C.S. All authors provided input to and approved the final version of the manuscript.

## Data archiving statement

Raw sequence data are available through the National Center for Biotechnology Information Sequence Read Archive under BioProject PRJNA722804.

## Competing interests

The authors declare no competing interests.

## Notes

### Competing Interest Statement

The authors have declared no competing interest.

