## Supplementary Figures for "The time course of molecular acclimation to seawater in a euryhaline fish"

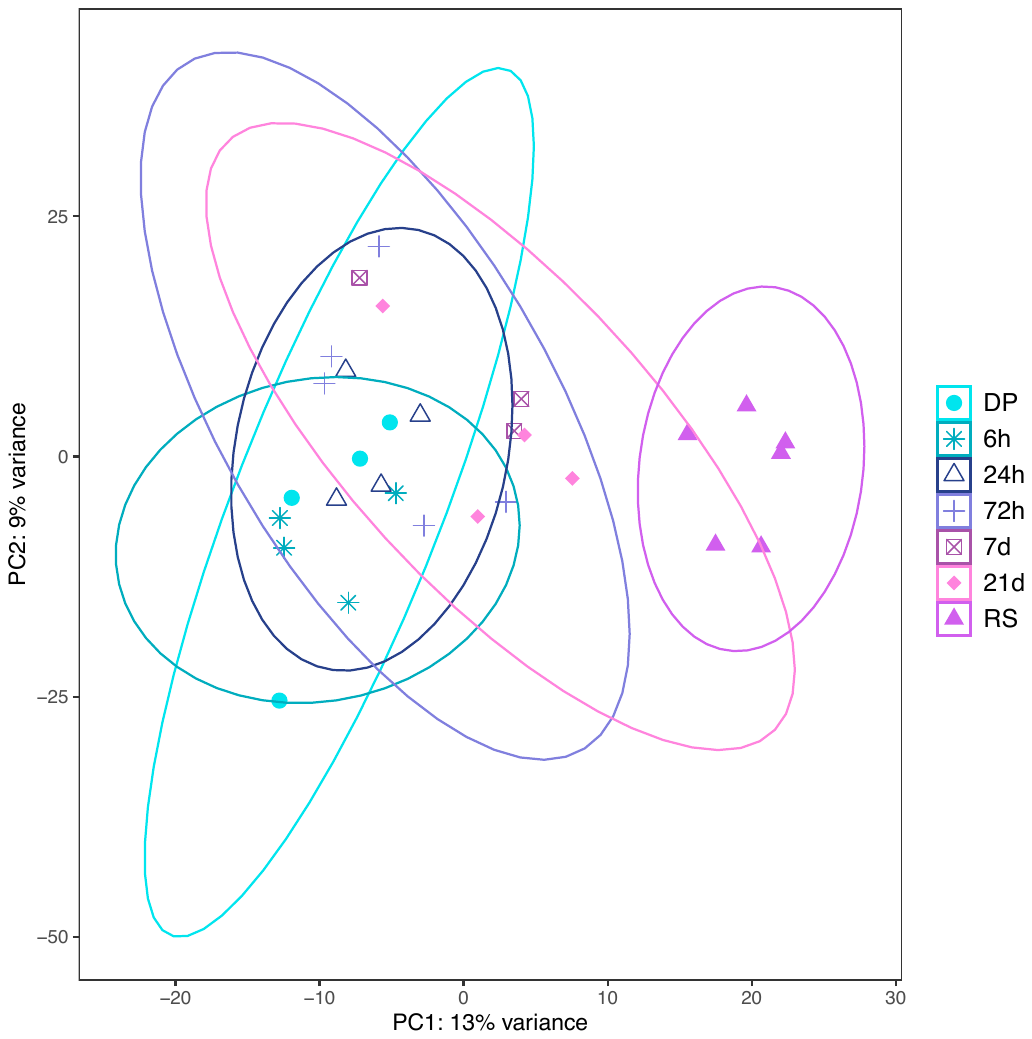

Supplementary Figure 1. Principal component analysis (PCA) of variance stabilized expression values for *Aphanius dispar* gills at the different sampling events. 22% of the total variation is explained by the first two components. Data ellipse for day 7 samples was not drawn because of the small sample size.

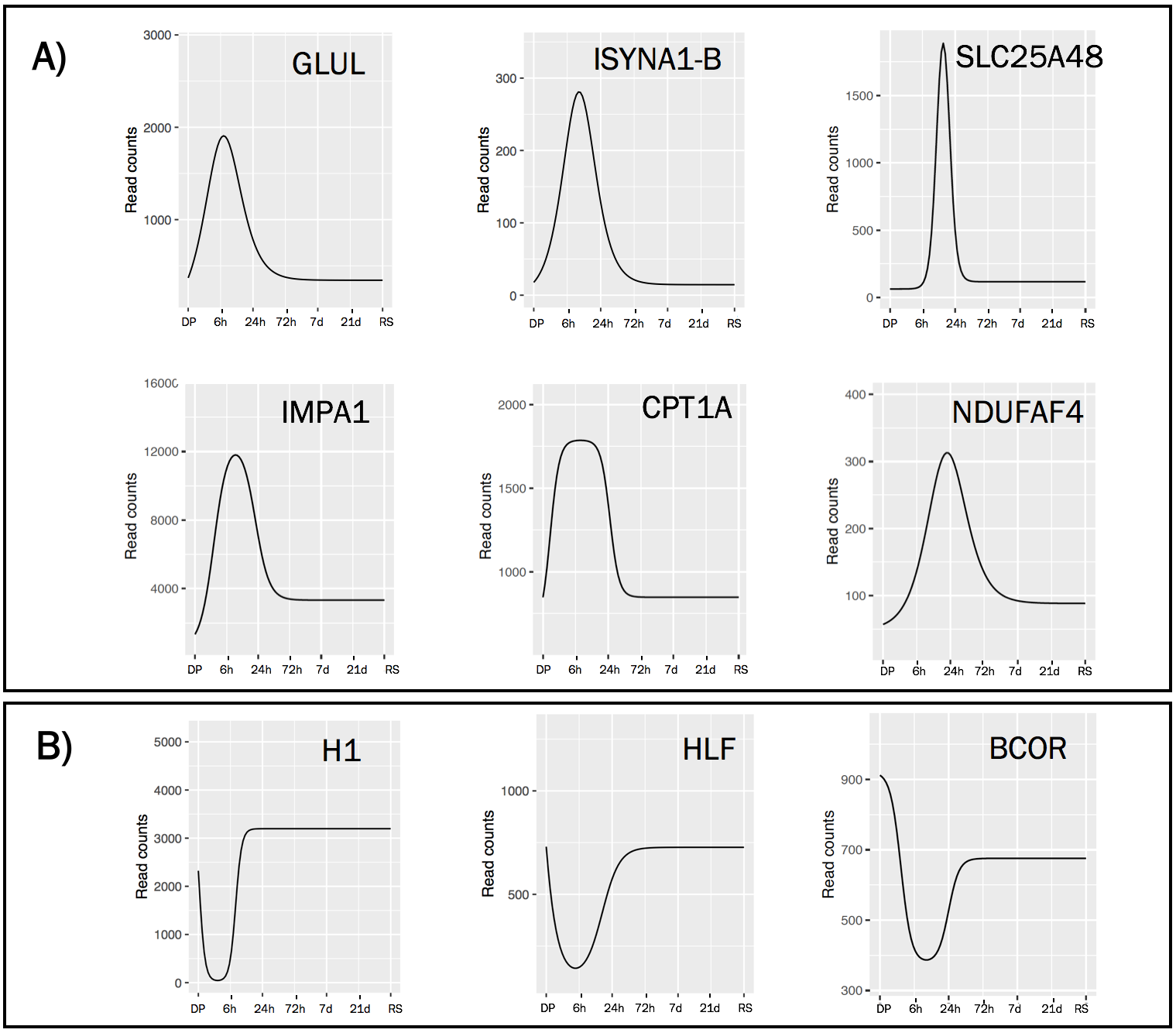

Supplementary Figure 2. Expression profiles of ImpulseDE2 identified transiently upregulated (A) and downregulated (B) genes in the first 24 hours.
